# A shared core microbiome in soda lakes separated by large distances

**DOI:** 10.1101/608471

**Authors:** Jackie K. Zorz, Christine Sharp, Manuel Kleiner, Paul M.K. Gordon, Richard T. Pon, Xiaoli Dong, Marc Strous

## Abstract

In alkaline soda lakes, high concentrations of dissolved carbonates establish an environment favouring productive phototrophic microbial mat communities. Here we show how different species of microbial phototrophs and autotrophs contribute to this exceptional productivity. Four years of amplicon and shotgun DNA sequencing data from microbial mats from four different lakes indicated the presence of over 2,000 different species of Bacteria and Eukaryotes. Metagenome-assembled-genomes were obtained for a core microbiome of <100 abundant bacteria, which was shared among lakes and accounted for half of the extracted DNA throughout the four year sampling period. Most of the associated species were related to similar microbes previously detected in sediments of Central Asian alkaline soda lakes, showing that common selection principles drive community assembly from a globally distributed reservoir of alkaliphile biodiversity. Dispersal events between the two distant lake systems were shown to be extremely rare, with dispersal rates a function of abundance in microbial mats, but not sediments. Detection of more than 7,000 expressed proteins showed how phototrophic populations allocated resources to specific processes and occupied complementary niches. Carbon fixation only proceeded by the Calvin-Benson-Bassham cycle, detected in Cyanobacteria, Alphaproteobacteria, and, suprisingly, Gemmatomonadetes. Our study not only provides new fundamental insight into soda lake ecology, but also provides a template, guiding future efforts to engineer robust and productive biotechnology for carbon dioxide conversion.

**Importance:** Alkaline soda lakes are among the most productive ecosystems worldwide, despite their high pH. This high productivity leads to growth of thick “mats” of filamentous cyanobacteria. Here, we show that such mats have very high biodiversity, but at the same time contain a core, shared set of only approximately 100 different bacteria that perform key functions, such as photosynthesis. This “core microbiome” occurs both in Canadian and Central Asian soda lakes, >8,000 km apart. We present evidence for (very rare) dispersion of some core microbiome members from Canadian mats to Central Asian soda lake sediments. The close similarity between distant microbial communities indicates that these communities share common design principles, that reproducibly lead to a high and robust productivity. We unravel a few examples of such principles and speculate that these might be applied to create robust biotechnology for carbon dioxide conversion, to mitigate of global climate change.

## Introduction

Soda lakes are among the most alkaline natural environments on earth, as well as among the most productive aquatic ecosystems known (1,2). The high productivity of soda lakes is due to a high bicarbonate concentration. Tens to hundreds of millimolars of bicarbonate are typically available for photosynthesis using carbon concentrating mechanisms (3,4), compared to generally < 2 mM in the oceans (5). This can lead to the formation of thick, macroscopic microbial mats with rich microbial biodiversity (6). Because of the high pH, alkalinity, and high sodium salinity of these environments, the microorganisms that reside in soda lakes are considered extremophiles (7). Using conditions of high pH and alkalinity is also a promising option to improve the cost-effectiveness of biotechnology for biological carbon dioxide capture and conversion (8-10).

Soda lakes have contributed to global primary productivity on a massive scale in Earth’s geological past (11). Currently, groups of much smaller soda lakes exist, for example, in the East African Rift Zone, rain-shadowed regions of California and Nevada, and the Kulunda steppe in South Russia (12). Many microorganisms have been isolated from these lakes. These include cyanobacteria (13-15), chemolithoautotrophic sulfide oxidizing bacteria (16-18), sulfate reducers (19,20), nitrifying (21-22) and denitrifying bacteria (23), as well as aerobic heterotrophic bacteria (24-25), methanotrophs (26), fermentative bacteria (27-28), and methanogens (29). Recently, almost one thousand Metagenome Assembled whole Genome sequences (MAGs) were obtained from sediments of Kulunda soda lakes (30).

In the present study we investigate the microbial mat community structure of four alkaline soda lakes located on the Cariboo Plateau in British Columbia, Canada. This region has noteworthy geology and biology due to the diversity in lake brine compositions within a relatively small region (31). There are several hundred shallow lakes on the Cariboo Plateau and these range in size, alkalinity, and salinity. Underlying basalt in some areas of the plateau, originating from volcanic activity during the Miocene and Pliocene eras, provides ideal conditions for forming soda lakes, as these areas are poor in calcium and magnesium (6,32,33). Some of these lakes harbor seasonal microbial mats that are either dominated by cyanobacteria or eukaryotic green algae. However, beyond this little is currently known about these systems in terms of microbiology.

We used a combination of shotgun metagenomes, and 16S and 18S rRNA amplicon sequencing to establish a microbial community structure for the microbial mats of four soda lakes. Next, we performed proteomics to show how specific populations allocate resources to specific metabolic pathways, focusing on photosynthesis, and carbon, nitrogen, and sulfur cycles. Overall, this study provides a comprehensive molecular characterization of a phototrophic microbial mat microbiome and shows how this highly productive ecosystem is supported by a set of complementary niches among phototrophs.

## Results and Discussion

The Cariboo Plateau contains hundreds of lakes of different size, alkalinity and salinity. Here we focused on four alkaline soda lakes (**Figure 1**) that feature calcifying microbial mats with similarities to ancient stromatolites or thrombolites (6,34,35). Between 2014 and 2017, the total alkalinity in these lakes was between 0.20-0.65 mol/L at pH 10.1-10.7 (**Supplementary Table 1**). Four years of amplicon sequencing data (16S and 18S rRNA) showed the microbial mats to be diverse communities, with 1,662 bacterial and 587 eukaryotic species-level operational taxonomic units (OTUs) identified, overall (**Supplementary Table 2**). The mat communities from different lakes were similar, but distinct, and relatively stable over time (**Figure 1**). Probe, Deer and Goodenough Lakes harbored predominantly cyanobacterial mats, whereas the mats of more saline Last Chance Lake contained mainly phototrophic Eukaryotes. This was shown with proteomics (see below), because it was impossible to compare abundances of Eukaryotes and Bacteria using amplicon sequencing. Bacterial species associated with 340 OTUs were found in all four lakes. These species accounted for 20.5% of the region’s species richness and 84% of the total sequenced reads, suggesting that there is a common and abundant “core” microbiome shared among the alkaline lakes of the Cariboo Plateau.

**Figure 1.**
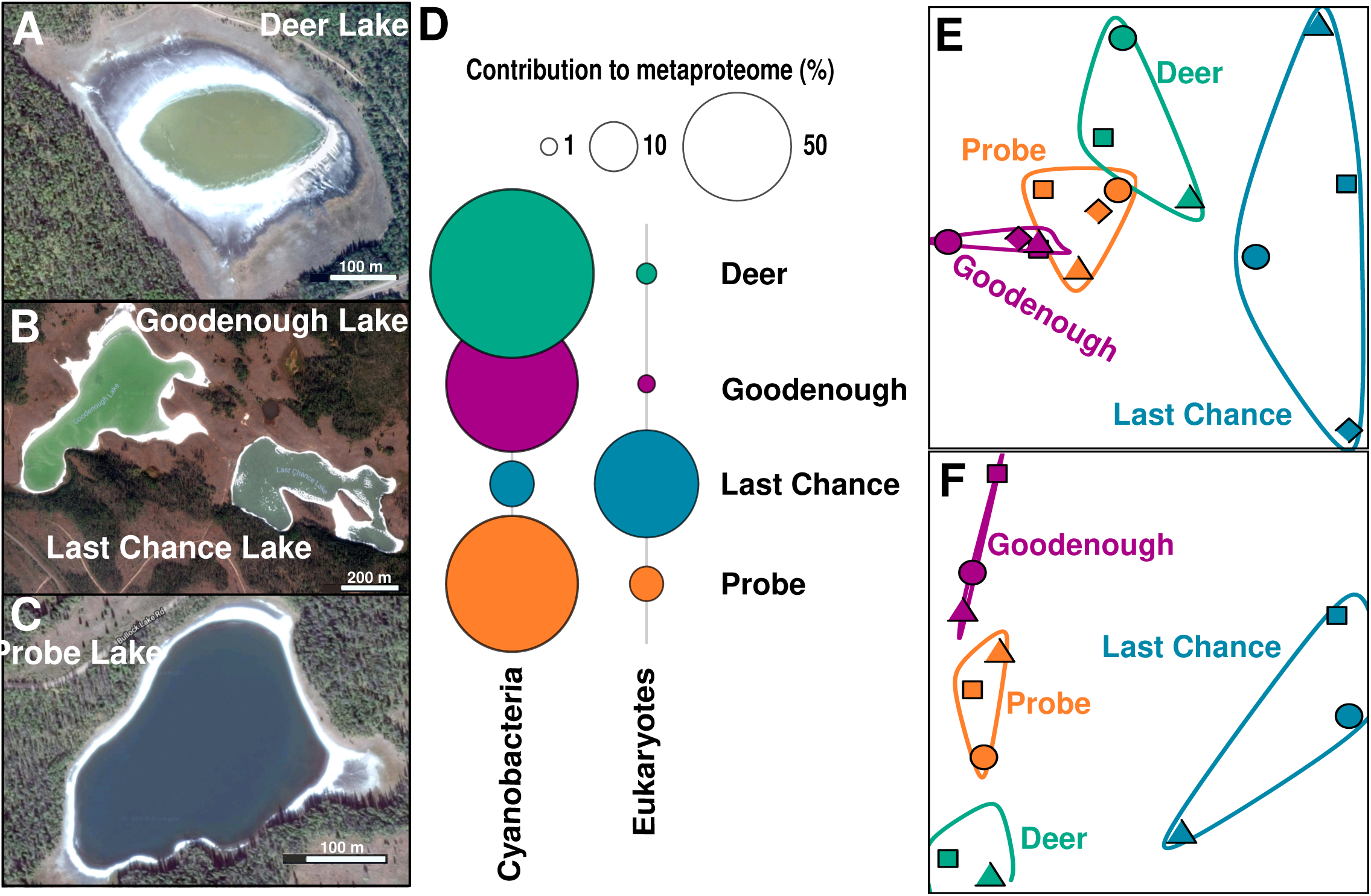
Satellite images of **A** Deer Lake, **B** Goodenough and Last Chance Lakes, **C** Probe Lake. **D.** Bubble plots showing the relative contribution of Cyanobacteria and Eukaryotes to the lake metaproteomes. **E.** Non-metric multidimensional scaling (NMDS) plots using Bray-Curtis dissimilarity to visualize the microbial communities of the soda lake mats over years of sampling using 16S rRNA amplicon sequencing data, and **F** 18S rRNA amplicon sequencing data. Shapes indicate year of sampling: Circles: 2014, square: 2015, diamond: 2016, triangle: 2017. Samples for 18S rRNA analysis were not taken in 2016, and Deer Lake samples were not taken in 2014 for 18S, and +2016 for 16S. NMDS Stress values were below 0.11.

After amplicon sequencing had outlined the core microbiome of the Cariboo soda lake microbial mats, shotgun metagenome sequencing, assembly and binning were used to obtain the provisional whole genome sequences, or metagenome-assembled genomes (MAGs), of its key microbiota. We selected 91 representative, de-replicated, near-complete (>90% for 85 MAGs), relatively uncontaminated (<5%, for 83 MAGs) for further analysis (**Supplementary Table 3**). For fifty-six MAGs, we independently assembled and binned 2-5 nearly identical (>95% average nucleotide identity) versions, indicating the presence of multiple closely related strains. 40-60% of quality-controlled reads were mapped to the 91 MAGs, showing that the associated bacteria accounted for approximately half of the DNA extracted. Most of the remaining reads were mapped to MAGs of lower quality and coverage, associated with a much larger group of less abundant bacteria. This was not surprising because amplicon sequencing had already indicated the presence of >2,000 different bacterial and eukaryotic species. Full length 16S rRNA gene sequences (**Supplementary Table 4**) were reconstructed from shotgun metagenome reads. Fifty-seven of those could be associated with a MAG based on taxonomic classifications and abundance profiles. Perfect alignment of full length 16S rDNA gene sequences to consensus OTU amplicon sequences showed that almost all these MAGs were core Cariboo microbiome members, present in each lake.

**Figure 2** shows the taxonomic affiliation and average relative sequence abundances for the bacteria associated with the MAGs. For taxonomic classification we used the recently established GTDB taxonomy (36). We also used the GTDB toolkit to investigate the similarity of the Cariboo mat genomes to >800 MAGs recently obtained from sediments of the Central Asian soda lakes of the Kulunda Steppe (30). The distance between the two systems of alkaline lakes is approximately 8,000 km. Yet, fifty-six of the Cariboo MAGs were clustered together with Kulunda MAGs and defined new family or genus level diversity in the context of the GTDB database (release 86, >22,000 whole genome sequences). This degree of similarity between geographically distant lake systems was surprising, especially because DNA was obtained from Kulunda sediments, not mats. It suggests that the core microbiome defined here for Cariboo lake mats, also applies to at least one other, well described system of soda lakes.

**Figure 2.**
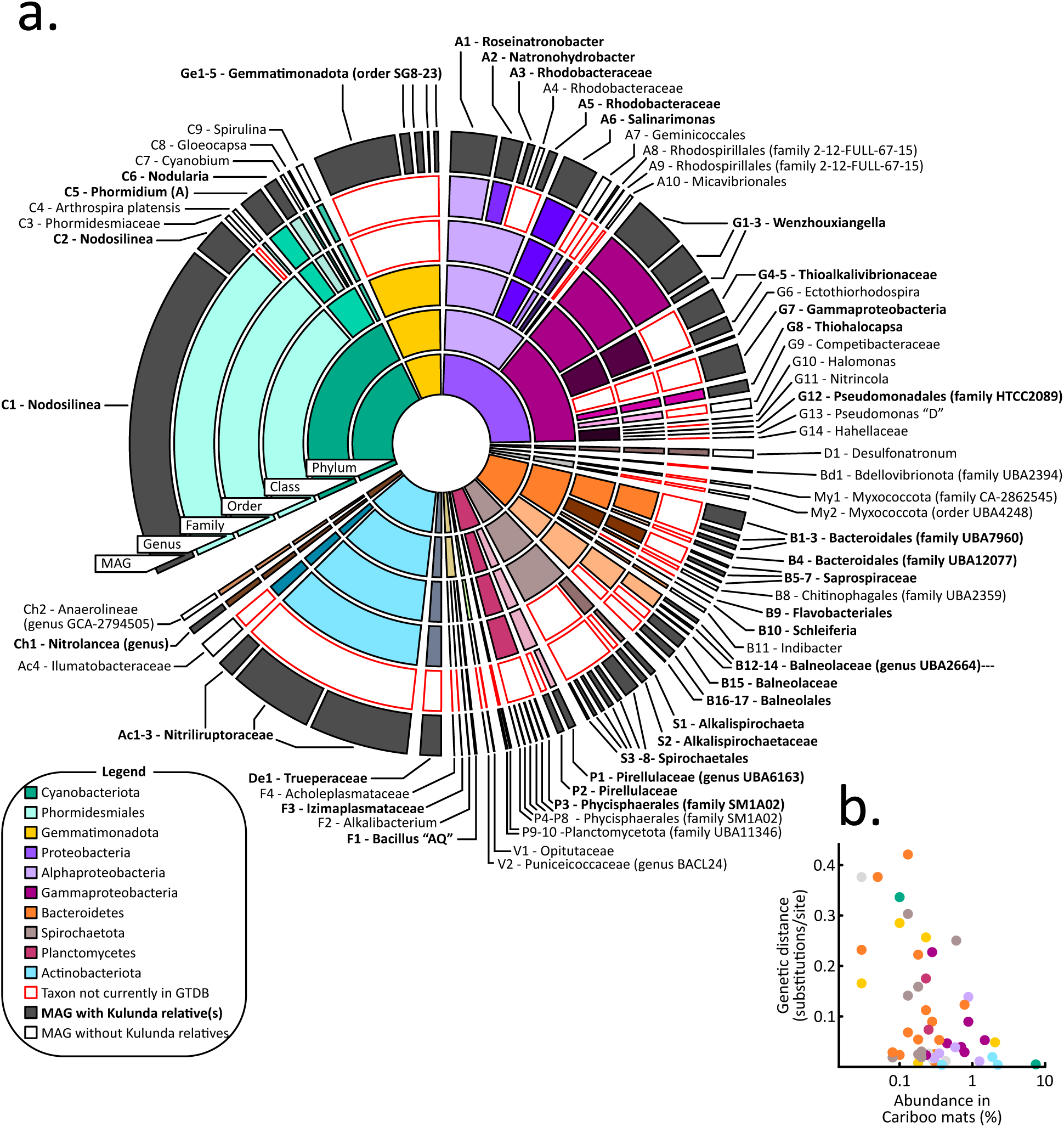
**a.** Sunburst diagram showing relative abundances and GTDB taxonomic classifications of metagenome-assembled-genomes (MAGs) obtained from Cariboo lakes. Core-microbiome MAGs with closest relatives among Central Asian (Kulunda) soda lake MAGs are shown in grey. Red outlines indicate new clades that were not yet represented in GTDB. For example, MAG C1, the most abundant MAG, is affiliated with the genus *Nodosilinea*, which was represented in GTDB, with a Kulunda MAG more similar than any genome present in GTDB. **b.** Scatter plot showing for each core microbiota the genetic distance between Cariboo and Kulunda representatives as a function of the abundance in Cariboo mat samples. This relationship is statistically significant (Pearson’s correlation r: -0.49, *p* < 0.05), but no such relationship was detected for the abundance of Kulunda MAGs. See also **Supplementary Table 3**.

Interestingly, the distance between the most similar MAGs from each of the two regions decreased with increasing abundance in Cariboo mats (Pearson correlation -0.49, *p* 0.0003, **Figure 2b, Supplementary Table 3**), but not with abundance in Kulunda sediments. For example, the most abundant Cariboo cyanobacterium (**C1** – affiliated with *Nodosilinea*, relative abundance >7%) displayed 99% average nucleotide identity over 85% of its genome with Kulunda MAG GCA_003550805. The latter displayed <0.1% relative abundance in Kulunda sediments. Mapping of Kulunda sequencing reads directly to Cariboo genomes (**Supplementary Table 3**) did not provide any evidence for the presence of previously undetected bacteria/MAGs in Kulunda sediments that were more similar to Cariboo bacteria/MAGs than those presented by Vavourakis et al. (2018).

These results suggest that when the Cariboo lakes formed ∼10,000 years ago after the last ice age (6), their microbiomes assembled from a much older, global reservoir of alkaliphile biodiversity. The striking relationship between Cariboo abundance and Kulunda-Cariboo relatedness might be explained by increased rates of successful dispersal/colonization for more abundant populations. Identification of vectors for dispersal still awaits future research, but bird migration is an obvious candidate. For example, the Northern Wheatear, which migrates between Northern Canada and Africa via Central Asia, could potentially link many known soda lakes worldwide. Abundance in sediments, located below mats, might not explain dispersal well, because sediments are less exposed to dispersal vectors than mats.

In any case, the genetic distances separating related bacteria were generally large, indicating that successful colonization by invading bacteria from a different lake system must be extremely rare. Possibly, only a single bacterium (MAG **C1**) traveled between and successfully colonized another lake system since the last ice age. A strong degree of isolation was also observed for other “ecological islands”, such as hot springs (37).

Thus, the observed similarities of the microbiota between distant lake systems indicate shared outcomes of community assembly for microbial mat microbiomes in two distant soda lake environments. Future studies will indicate whether the core microbiota of Kulunda and Cariboo soda lakes has also assembled in other soda lakes.

Dispersal between Cariboo soda lakes, separated by at most 40 km, was very effective. For all 56 sets of 2-5 nearly identical MAG variants (average nucleotide identity >95%) we detected co-occurrence of all variants (**Supplementary Table 5**). This also showed that competitive exclusion was irrelevant, even for these nearly identical bacteria. Comparison of ratios of synonymous and non-synonymous mutations among the most rapidly evolving core genes – genes present in all genome variants, **Supplementary Table 6** – showed that diversifying selection acted on 775 genes, including many transporters and genes involved in cell envelope biogenesis. Accessory genes – not encoded on all variant genomes – and CRISPRs could display many more ecologically relevant differences, which could prevent competitive exclusion.

The processes that dictate assembly of effective phototrophic microbial mat communities are well understood, with ecological adaptations and responses to dynamic light, oxygen, sulfide, pH and carbon dioxide gradients (38). But, to what extent do these known “rules of engagement” also apply to alkaline soda lake microbial mats, where primary productivity has access to unlimited inorganic carbon (2,6)? We performed environmental proteomics and connected protein expression to abundant MAGs to answer this question for the Cariboo Plateau soda lake mats (**Supplementary Table 7**).

Over seven thousand expressed proteins were identified, with high confidence, in daytime mat samples from each of the lakes. For comparison, the most comprehensive environmental proteomes obtained so far have identified up to approximately ten thousand proteins (39). Given the high diversity and extremely complex nature of the mat samples, identification of 7,217 proteins is an excellent starting point for ecophysiological interpretation. Approximately half of the expressed proteins could be attributed to the 91 MAGs, consistent with abundance estimates inferred from amplicon and shotgun data. This enabled us to investigate how the bacteria associated with the MAGs distributed their resources over different ecophysiological priorities (40). Given that a substantial amount of cellular energy goes towards manufacturing proteins, the relative proportion of a proteome dedicated to a particular function provides an estimate of how important that function is to the organism. Proteomic data were also used to estimate the ^13^C content of some abundant species, providing additional information on which carbon source they used and to what extent their growth was limited by carbon availability (Kleiner et al., 2018). Brady et al. (2013) previously showed that microbial mat organic matter had δ^13^C values of -19 to -25‰, up to 11.6‰ depleted in ^13^C compared to bulk inorganic carbonates, consistent with non-CO_2_-limited photosynthesis. Overall protein δ^13^C values for the four lakes inferred from the proteomics data in the present study were between -19 and -25‰, consistent with previous results for mat organic matter.

Consistent with their reputation as productive ecosystems with virtually unlimited access to inorganic carbon, the most abundant bacteria were large, mat-forming (filamentous) cyanobacteria, related to *Nodosilinea* and *Phormidium*. Pigment antenna proteins and photosynthetic reaction center proteins accounted for the largest fraction of detected proteins overall. The organism with the highest presence in the metaproteome was the cyanobacterial MAG **C1**, affliated with *Nodosilinea* and accounting for up to 42% of mat metaproteomes. Remarkably, we were able to identify 1,103 proteins from this MAG, 27% of its predicted proteome (**Figure 3**). This level of detection is comparable to results of pure cultures of cyanobacteria, such as *Arthrospira, 21%, and Cyanothece, 47%* (41,42). Nine cyanobacterial MAGs were assembled in total, and proteins from all nine were detected in the metaproteomes of all four lakes (**Supplementary Table 7**). It is clear that the presence of so many cyanobacteria provides functional redundancy and contributes to functional robustness and resiliency (43,44). However, we also detected strong evidence for niche differentiation for those cyanobacteria with larger numbers of proteins detected, in particular MAG **C1** (*Nodosilinea*), and MAG **C5** (*Phormidium “A”*) (**Figure 4**).

**Figure 3.**
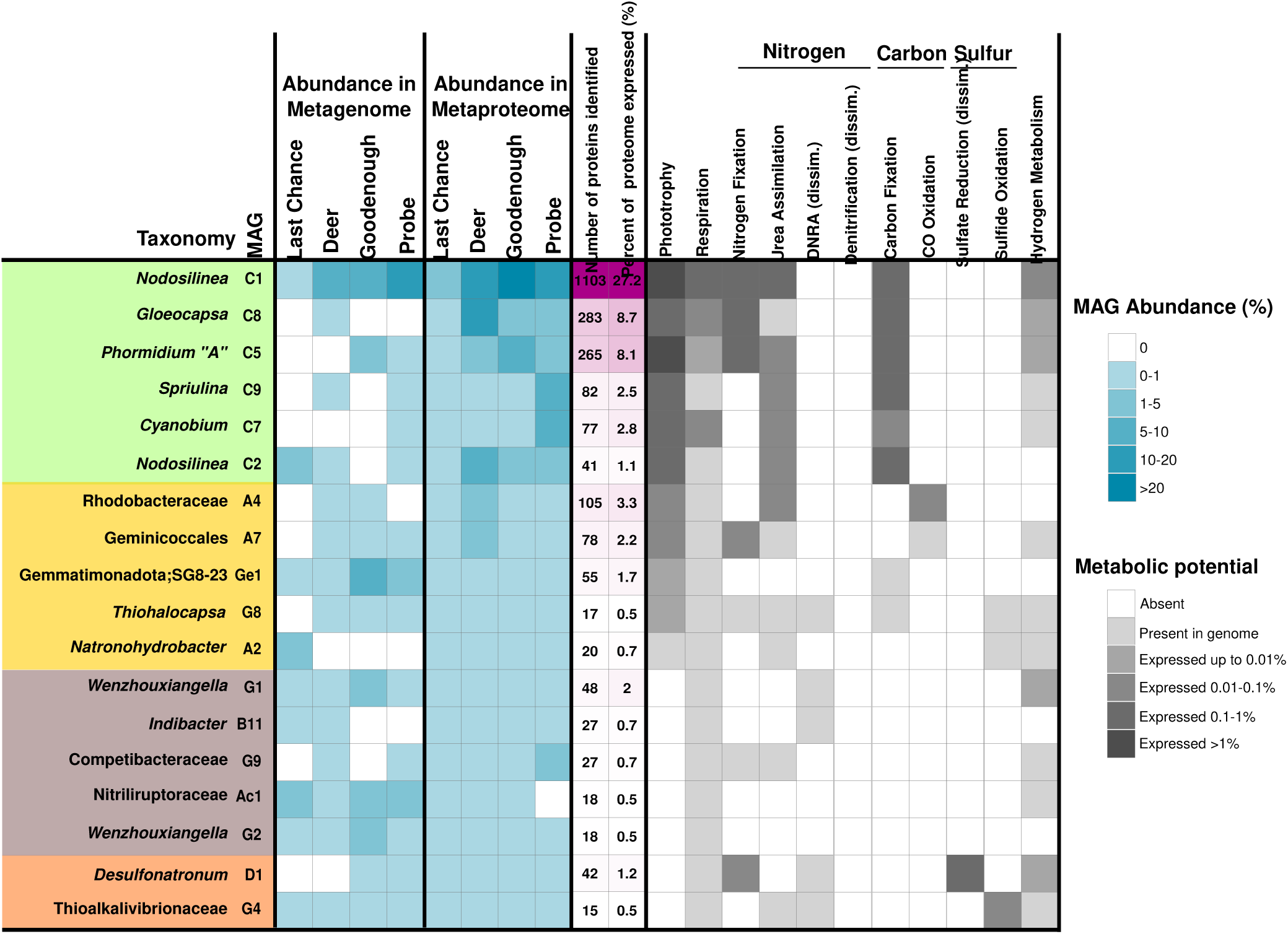
Heatmap showing abundances and expressed functions for metagenome-assembled genomes (MAGs) with at least 15 proteins identified in the metaproteomes. MAGs are broadly arranged based on function, with photoautotrophs in green, anoxygenic phototrophs in yellow, sulfur cycling in orange, and other heterotrophic bacteria in brown. Metabolic potential was inferred from the genes listed in **Supplementary Table 7**. If the gene was identified in a metaproteome it was considered “expressed”, and is shaded according to its highest relative abundance (% of all peptide spectral matches) in the four lake metaproteomes.

**Figure 4.**
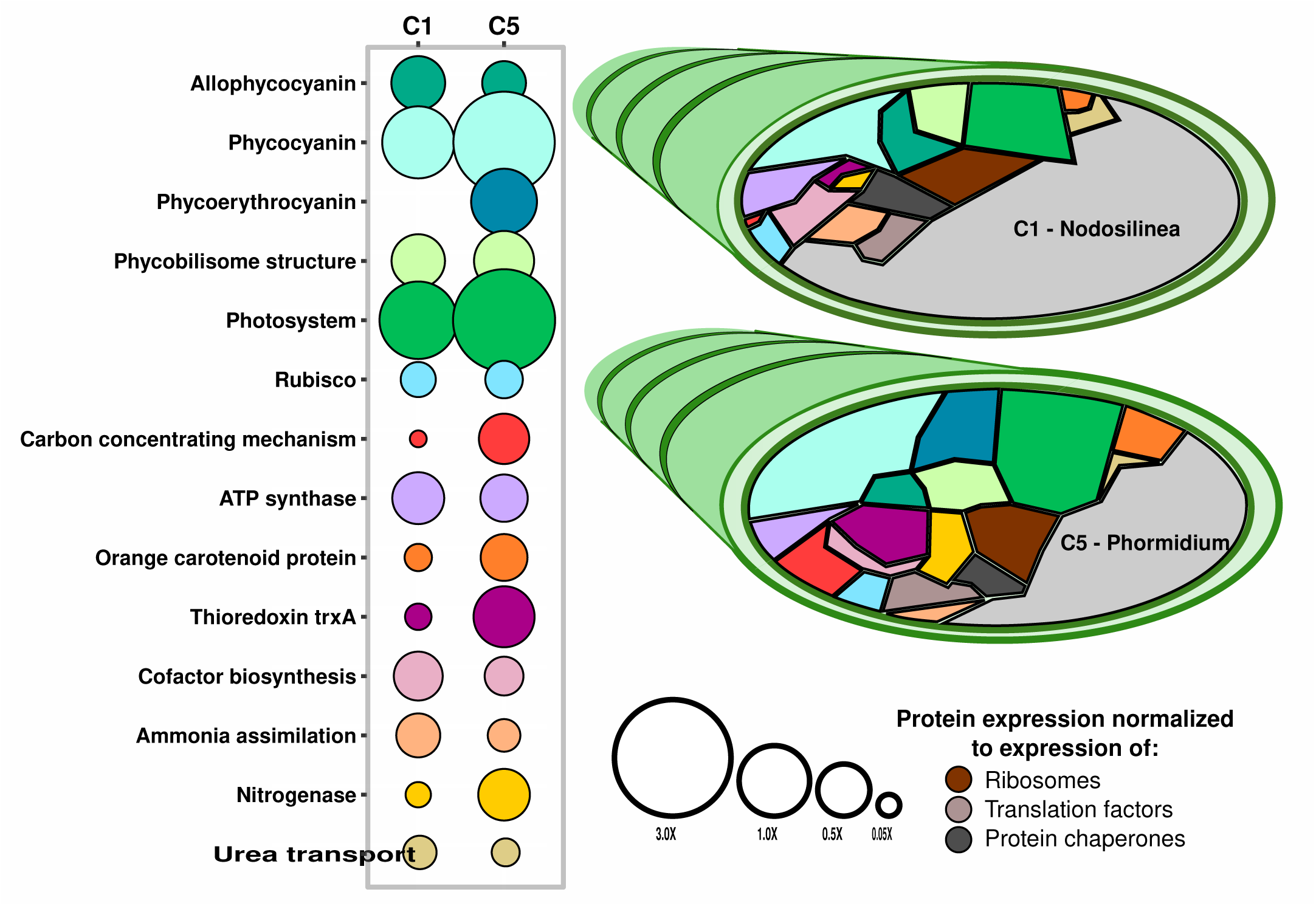
Voronoi diagrams comparing expression levels of functions by MAGs **C1** and **C5**, both associated with filamentous cyanobacteria. The area of for each function is proportional to the percent that protein or subsystem accounts for out of the MAG’s expressed proteins. Size of the bubble in the bubble plot is normalized against the relative abundances of ribosomal proteins, translation factors, and protein chaperones in the MAG’s proteome. See also **Supplementary Table 7**.

Phycobilisomes, the large, proteinaceous, light harvesting complexes of cyanobacteria, contain an assortment of pigments which absorb at different wavelengths of light, and re-emit that light at longer wavelengths, around 680 nm, compatible with the reaction center of Photosystem II. Phycobilisome pigment composition varied among the cyanobacterial populations, leading to niche differentiation based on light quality, as was also observed in the marine environment (45). **C1** and most other cyanobacterial populations expressed high amounts of phycocyanin, maximum absorbance 620 nm, and allophycocyanin, maximum absorbance 650 nm. In contrast, **C5** uniquely expressed the pigment phycoerythrocyanin, with a maximum absorbance at 575 nm (**Figure 4**). Phycoerythrocyanin would enable this population to absorb shorter wavelengths of light, in comparison to its cyanobacterial neighbours, and expands the “spectral reach” of photosynthesis for these mat communities, increasing productivity. The absence of expression of phycoerythrin, which has a maximum absorbance at 495 and 560 nm, is consistent with the light attenuation profile of aquatic environments with high dissolved organic matter, such as productive alkaline lakes, where wavelengths < 500 nm are rapidly attenuated (46,47).

Shorter wavelength light (blue/green light) has higher energy, and high energy photons can damage photosynthetic machinery in cyanobacteria. If **C5** would be exposed to these photons, as its pigment profile suggests, this could lead to more photodamage. Consistently, this population displayed higher expression of proteins like thioredoxin, for scavenging reactive oxygen species, and orange carotenoid protein for photoprotection (**Figure 4**).

Inorganic carbon fixation and acquisition are central to realizing high primary productivity and the associated enzymes were highly expressed. The rate-limiting, Calvin-Benson-Bassham Cycle (CBB) enzyme RuBisCO accounted for approximately 1% of the expressed proteomes of cyanobacterial MAGs, large fraction for a single enzyme (**Figure 4**). In contrast, the expression of the carbon concentrating mechanism (CCM, needed for bicarbonate uptake) varied greatly among cyanobacteria. In **C1** and **C8**, CCM proteins accounted for less than 0.2% of the proteomes. In **C5**, CCM proteins accounted for almost 3% of the expressed proteomes. **C5** was the only population to express CCM proteins to a greater level than RuBisCO proteins, suggesting that this population’s growth rate might be limited by bicarbonate availability. Indeed, **C5**’s δ^13^C value was -20.6±2.7‰, compared to -25.2±0.8‰ for **C1**. A decrease in isotopic fractionation during photosynthesis is usually associated with CO_2_(or bicarbonate) limitation (48). We might conclude that **C5**’s access to higher energy radiation leads to a higher rate of photosynthesis, increased oxygen production, a higher need for protection against free radicals, a higher growth rate against a limiting rate of bicarbonate supply. At a relative abundance of up to 2.3%, **C5** was not the most abundant cyanobacterium, so if it had a higher growth rate, it must also have had a higher decay rate, which is typical for this organism appearing to be an ecological R strategist.

Nitrogen is a commonly limiting nutrient for primary production in soda lakes globally (49). The Cariboo Plateau lakes also display low or undetectable concentrations of ammonium and nitrate in lake waters (**Supplementary Table 1**). Consistently, no expression was detected for any proteins involved in nitrogen loss processes, such as nitrification or denitrification, or for assimilatory nitrate reductases or nitrate transporters.

Many bacteria, including the cyanobacteria **C1, C5** and **C8**, expressed the key genes for the energetically expensive process of nitrogen fixation (**Supplementary Table 7**). All cyanobacteria further expressed glutamine synthetase, for the assimilation of ammonia under nitrogen limiting conditions (50), and the urea transporter. Dinitrogen, urea and, possibly, ammonia, were apparently the main nitrogen sources supporting photosynthesis. Parallel performance of nitrogen fixation by different bacteria provided functional redundancy, contributing to functional robustness and resiliency.

Phosphate can also be a limiting nutrient in soda lakes (49), and this appeared to be the case for Deer Lake in the present study, where phosphate was undetectable in lake waters (**Supplementary Table 1**). Cyanobacterium **C8** (*Gloeocapsa*) was the most abundant population in Deer Lake (12.9% of Deer Lake metaproteome), and expressed a high-affinity phosphate transport system at higher levels (1.5% of C8 expressed proteome) than the other cyanobacteria. Phosphate potentially limited primary production in Deer Lake, as anoxygenic photoheterotrophs were 4-40x more abundant here than in the other lakes (**Figure 3, Supplementary Tables 3** and **7**).

The microbial mats of the Cariboo region display steep oxygen and sulfide gradients (6), providing opportunities for photoheterotrophic bacteria that use any remaining light, which penetrates beyond the oxic layer created by cyanobacteria (38,51). Photosystem proteins such as Puf or Puh were expressed by purple non-sulfur bacteria affiliated with Rhodobacteraceae, MAG **A4**, and Geminicoccales, MAG **A7**, as well as autotrophic purple sulfur bacteria, affiliated with *Thiohalocapsa*, MAG **G8**. Both photoheterotrophs were relatively abundant in phosphate-limited Deer Lake, at 3.2% and 2.8% respectively. In addition to PuhA, MAG **A4** expressed all three subunits of carbon monoxide dehydrogenase (coxSML). Carbon monoxide could be produced by photooxidation of organic material (52), and could serve as an alternative energy source for these bacteria. Organic substrates supporting photoheterotrophic growth likely consist of cyanobacterial fermentation products, glycolate from photorespiration (38) or could originate from biomass decay. By re-assimilation of organic matter or re-fixation of bicarbonate using light energy, these organisms enhance the overall productivity of the mats.

Most unexpected among photoheterotrophs was population **Ge1**, a representative of an uncultured family within the recently defined phylum Gemmatimonadota. This particular population expressed the PufC subunit of the photosynthetic reaction center and contains the remaining photosystem genes in its genome (PufLMA, PuhA, AcsF). The ability for members of this phylum to use light energy was only recently discovered (53), and the capacity for phototrophy appears to be widespread among members of that phylum (54).

The Gemmatimonadetes bacterium isolated by Zheng and colleagues is heterotrophic, without evidence for a carbon fixation pathway. Interestingly, MAG **Ge1** is in possession of all the genes required for a complete carbon-fixing CBB cycle. Genes homologous to the functional RuBisCO Form 1C large subunit (RbcL), RuBisCO small subunit (RbcS) were identified, as well as a copy of the CBB cycle-specific enzyme Phosphoribulokinase (PRK). These genes were arranged sequentially in the genome: RbcS, RbcL, and PRK, an arrangement that points at facultative autotrophy (55). Upon further investigation of the published MAGs from the Kulunda Steppe soda lakes in Central Asia, we found five additional Gemmatimonadetes MAGs (**Figure 5**), that encoded these three CBB cycle genes with the same synteny, and with 88-98% amino acid identity, to the genes of **Ge1**. All identified RbcL genes are functional Form 1C RbcL sequences (**Figure 5B**). To our knowledge these six MAGs contain the first examples of the full suite of CBB cycle genes in this phylum. Given the large number of amino acids (>90%) shared with homologuous genes encoded in Alphaproteobacteria (e.g. Rhizobiales bacterium YIM 77505 RbcL), it seems likely that the last common ancestor of these Gemmatimonadetes populations acquired the CBB genes via horizontal gene transfer from an Alphaproteobacterium, prior to the dispersal and speciation of the clade into the Kulunda Steppe and Cariboo Plateau populations. We did not detect expression for these genes and were not able to estimate the δ^13^C value for this bacterium (too few high quality MS1 spectra) so it remains unknown to what extent this bacterium used bicarbonate as a carbon source.

**Figure 5.**
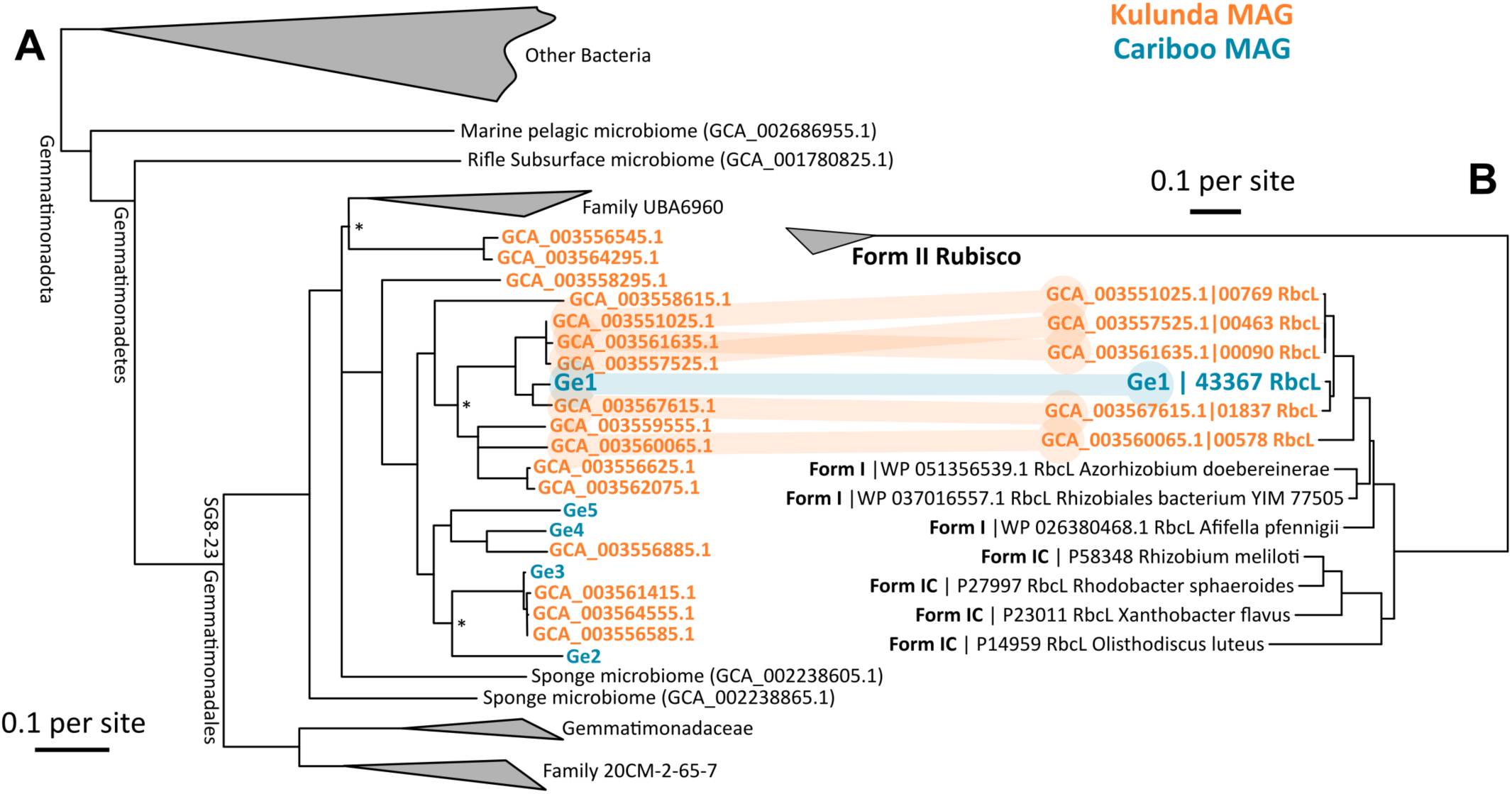
**a.** Phylogenetic tree of MAGs affiliated with Gemmatimonadota obtained from Cariboo lakes (blue, Ge1-5) and Kulunda lakes (orange). The tree was created with GTDBtk, based on concatenated conserved single copy genes, using fasttree2. b. Phyloegentic tree of the RuBisCO Form 1 encoded on MAGs in one of the Gemmatimonadota clades. Congruence between the trees indicates vertical inheritance after a single horizontal gene transfer event from Alphaproteobacteria.

The presence of the autotrophic purple sulfur bacterium **G8**, affiliated with *Thiohalocapsa*, indicated active sulfur cycling within the mats, as expected based on sulfide gradients (Brady et al., 2013). Indeed, MAG **D1**, affiliated with *Desulfonatronum* (20,56) expressed *aprAB, sat*, and *dsrAB*. It also expressed an alcohol dehydrogenase, a formate dehydrogenase, and a hydrogenase, indicating that it oxidized compounds such as ethanol, formate, and hydrogen. These could be derived from dark fermentation by cyanobacteria or from decaying biomass. Sulfide produced by **D1** was likely re-used by MAGs **G8** and **G4**, the latter affiliated with *Thioalkalivibrionaceae* (18,57). **G4** expressed *soxX, soxC, dsrA*, and *fccB*, suggesting sulfide oxidation through both the sox pathway and the reverse dsr pathway. Expression of *sox* and *fcc* was also detected for other unbinned populations, affiliated with Alphaproteobacteria, Chromatiales, and other Gammaproteobacteria.

In conclusion, we used metaproteomes and metagenomes to address fundamental questions on the microbial ecology of soda lake mats. We obtained 91 metagenome assembled genomes and showed that part of these taxa define a core microbiome, a group of abundant bacteria present in all samples over space (four lakes) and time (four years). We showed that a very similar community assembled independently in Central Asian soda lakes. The similarity between some of the microbial genomes found in these soda lake regions, incredible in the light of their vast physical separation, suggests that vectors for dispersal are generally ineffective, but can sometimes distribute abundant community members at the global scale. We also showed both functional redundancy and existence of complemental niches among cyanobacteria, with evidence for K and R strategists living side by side. The nature and origin of carbon sources for photoheterotrophs, including potentially mixotrophic Gemmatimonadetes is an exciting avenue for future research. The presented core microbiome provides a blueprint for design of a productive and robust microbial ecosystem that could guide effective biotechnology for carbon dioxide conversion.

## Materials and Methods

### Study Site and Sample collection

Samples from benthic microbial mats were collected from four lakes in the Cariboo Plateau region of British Columbia, Canada in May of 2014, 2015, 2016, and 2017. Microbial mats from Last Chance Lake, Probe Lake, Deer Lake, and Goodenough Lake were sampled (coordinates in **Supplementary Table 1**). Mats were immediately frozen, transported on dry ice, and stored at-80°C within 2 days of sampling. In 2015 and 2017, water samples for aqueous geochemistry were also taken and stored at -80°C until analysis.

### Aqueous Geochemistry

Frozen lake water samples were thawed and filtered through a 0.45 µm nitrocellulose filter (Millipore Corporation, Burlington, MA) prior to analysis. Carbonate/bicarbonate (HCO_3_ ^-^) alkalinity analysis was conducted using an Orion 960 Titrator (Thermo Fisher Scientific, Waltham, MA), and concentrations were calculated via double differentiation using EZ 960 software. Major cations (Ca^2+^, Mg^2+^, K^+^, and Na^+^) were analyzed using a Varian 725-ES Inductively Coupled Plasma Optical Emission Spectrophotometer (ICP-OES). Major anions (Cl^-^, NO_3_^-^ and SO_4_^2-^) were analyzed using a Dionex ICS 2000 ion chromatograph (Dionex Corporation, Sunnyvale, CA), with an Ion Pac AS18 anion column (Dionex Corporation, Sunnyvale, CA).

Water for reduced nitrogen quantification was filtered through a 0.2 µm filter (Pall Life Sciences, Port Washington, NY). Concentrations were measured using the ortho-phthaldialdehyde fluorescence assay as previously described (58), with excitation at 410 nm, and emission at 470 nm.

### Amplicon sequencing and data processing

DNA extraction and amplicon sequencing were performed as previously described (10), with primer sets TAReuk454FWD (565f CCAGCASCYGCGGTAATTCC) and TAReukREV3 (964b ACTTTCGTTCTTGATYRA), targeting Eukaryota, and S-D440 Bact-0341-a-S-17 (b341, TCGTCGGCAGCGTCAGATGTGTATAAGAGACAGCCTACGGGAGGCAGCAG), and S-D-Bact-0785-a-A-21 (805R, GTCTCGTGGGCTCGGAGATGTGTATAAGAGACAGGACTA CHVGGGTATCTAATCC) targeting Bacteria. Sequencing was performed using the MiSeq Personal Sequencer (Illumina, San Diego, CA) using the 2 x 300 bp MiSeq Reagent Kit v3. The reads were processed with MetaAmp (59). After merging of paired end reads (>100bp overlap and <8 mismatches in the overlapping region), primer trimming and quality filtering (<2 mismatches in primer regions and at most 1 expected error), trimming to 350bp, reads were clustered into operational taxonomic units (OTUs) of >97% sequence identity. Statistics (ANOSIM, Mantel correlations using conductivity, anion and cation concentrations) and visualization (non-metric multidimensional scaling, NMDS) were performed in R, using *vegan* (60). For NMDS, OTUs <1% abundant in all samples were excluded, as were those affiliated with Metazoa, because of large variations in rRNA copy and cell numbers.

### Shotgun metagenome sequencing and data processing

Metagenomes of 2015 mat samples were sequenced as described previously (61). Briefly, DNA was sheared into fragments of ∼300 bp using a S2 focused-ultrasonicator (Covaris, Woburn, MA). Libraries were created using the NEBNext Ultra DNA Library Prep Kit (New England Biolabs, Ipswich, MA) according to the manufacturer’s protocol, which included a size selection step with SPRIselect magnetic beads (Beckman Coulter, Indianapolis, IN) and PCR enrichment (8 cycles) with NEBNext Multiplex Oligos for Illumina (New England Biolabs, Ipswich, MA). DNA concentrations were estimated using qPCR and the Kapa Library Quant Kit (Kapa Biosystems, Wilmington, MA) for Illumina. 1.8 pM of DNA solution was sequenced on an Illumina NextSeq 500 sequencer (Illumina, San Diego, CA) using a 300 cycle (2 x 150 bp) high-output sequencing kit at the Center for Health Genomics and Informatics in the Cumming School of Medicine, University of Calgary. Raw, paired-end Illumina reads were filtered for quality as previously described (62). After that, the reads were coverage-normalized with BBnorm (sourceforge.net/projects/bbmap) with “target=100 min=4”. Overlapping reads were merged with BBMerge with default settings. All remaining reads were assembled separately for each library with MetaSpades version 3.10.0 (63), with default parameters. Contigs of <500 bp were not further considered. tRNA, ribosomal RNA, CRISPR elements, and protein-coding genes were predicted and annotated using MetaErg (sourceforge.net/projects/metaerg/). Per-contig sequencing coverage was estimated and tabulated by read mapping with BBMap, with default settings and “jgi_summarize_bam_contig_depths”, provided with MetaBat (64). Each assembly was binned into Metagenome-Assembled-Genomes (MAGs) with MetaBat with options “-a depth.txt –saveTNF saved_2500.tnf –saveDistance saved_2500.dist -v –superspecific -B 20 --keep”. MAG contamination and completeness was estimated with CheckM (65). MAGs were classified with GTDBtk (version 0.2.2, database release 86) (36), together with MAGs previously obtained from Kulunda soda lakes (30). fastANI was used to compare MAGs across libraries/assemblies (66). Relative sequence abundances of MAGs were estimated based on contig sequencing coverage. 16S rRNA gene sequences were obtained with Phyloflash2 (67) and were associated with MAGs based on phylogeny and sequencing coverage covariance across samples, and to OTUs based on sequence identity. The RuBisCO phylogenetic tree was created with MEGA (68). Core genes of MAG variants were identified using blast and these genes were used to determine the abundances of variants across samples using BBMap, with parameters minratio=0.9 maxindel=3 bwr=0.16 bw=12 fast ambiguous=toss. To identify diversified core genes, variants were aligned with mafft (69) and only genes with >50 single nucleotide polymorphisms (SNPs), >1% of positions with a SNP, and with a fraction of non-synonymous SNPs of >0.825 were kept.

### Protein Extraction and metaproteomics

Protein was extracted and analyzed from 2014 mat samples, as previously described (61). Briefly, lysing matrix bead tubes A (MP Biomedicals) containing mat samples and SDT-lysis buffer (0.1 M DTT) in a 10:1 ratio were bead-beated in an OMNI Bead Ruptor 24 for 45 seconds at 6 m/s. Next, tubes were incubated at 95°C for 10 minutes, spun down for 5 min at 21,000 *g* and tryptic peptides were isolated from pellets by filter-aided sample preparation (FASP) (70). Peptides were separated on a 50 cm × 75 µm analytical EASY-Spray column using an EASY-nLC 1000 Liquid Chromatograph (Thermo Fisher Scientific, Waltham, MA) and eluting peptides were analyzed in a QExactive Plus hybrid quadrupole-Orbitrap mass spectrometer (Thermo Fisher Scientific). Each sample was run in technical quadruplicates, with one quadruplicate run for 260 minutes with 1 µg of peptide loaded, and the other three for 460 minutes each, with 2-4 µg of peptide loaded.

Expressed proteins were identified and quantified with Proteome Discoverer version 2.0.0.802 (Thermo Fisher Scientific), using the Sequest HT node. The Percolator Node (71) and FidoCT were used to estimate false discovery rates (FDR) at the peptide and protein level respectively. Peptides and proteins with DFR >5% were discarded. Likewise, proteins without protein-unique-peptides, or <2 unique peptides were discarded. Relative protein abundances were estimated based on normalized spectral abundances (72). The identification database was created using predicted protein sequences of binned and unbinned contigs, after filtering out highly similar proteins (>95% amino acid identity) with cd-hit (73), while preferentially keeping proteins from binned contigs. Sequences of common contaminating proteins were added to the final database (http://www.thegpm.org/crap/), which is available under identifier PXD011230 in ProteomeXchange. In total, 3,014,494 MS/MS spectra were acquired, yielding 298,187 peptide spectral matches, and 7,217 identified proteins.

### Data availability

Amplicon sequences can be found under the Bioproject PRJNA377096. The 16S rRNA sequence Biosamples are: SAMN06456834, SAMN06456843, SAMN06456852, SAMN06456861, SAMN09986741-SAMN09986751, and the 18S rRNA sequence Biosamples are: SAMN09991649-SAMN09991660. The metagenome raw reads and metagenome assembled genomes can also be found under the Bioproject PRJNA377096. The Biosamples for the metagenome raw reads are SAMN10093821-SAMN10093824, and the Biosamples for the MAGs are SAMN10237340-SAMN10237430. The metaproteomics data has been deposited to the ProteomeXchange Consortium via the PRIDE partner repository (74) with the dataset identifier PXD011230.

## Acknowledgements

We thank the University of Calgary’s Center for Health Genomics and Informatics for sequencing and informatics services. We thank Michael Nightingale and Agasteswar Vadlamani for help with analysis of aqueous geochemistry. We also thank Timber Gillis, Hayely Todesco, Harsimrit Lakhyan, and Sydney Urschel for help with sample collection and DNA extractions. We would like to thank Dan Liu and Angela Kouris for help with metaproteomics sample preparation and analysis. We thank Carmen Li for help with MiSeq sequencing, and Maryam Ataeian for help with metagenome analysis. This study was supported by the Natural Sciences and Engineering Research Council (NSERC), Canada Foundation for Innovation (CFI), Canada First Research Excellence Fund (CFREF), Genome Canada, Western Economic Diversification, the International Microbiome Center (Calgary), Alberta Innovates, the Government of Alberta, and the University of Calgary.

## Supplementary Tables available as 10.6084/m9.figshare.7991171

Table 1 – Aqueous Geochemistry of the four lakes.

Table 2 – Operational Taxonomic Units, Bacterial 16S and 18S.

Table 3 – Metagenome Assembled Genomes (MAGs) – GTDB classification, abundances, quality, relationships to Kulunda MAGs.

Table 4 – Full length 16S rRNA gene sequences associated with MAGs.

Table 5 – Co-occurrences of nearly identical variants of MAGs, showing no evidence for competitive exclusion.

Table 6 – Evidence for diversifying evolution among some core genes of sets of MAG variants.

Table 7 – Expression data for signature genes of different metabolic pathways (Figure 4).

